# On the ecology and distribution of steelhead *(Oncorhynchus mykiss)* in California’s Eel River

**DOI:** 10.1101/2020.03.18.996934

**Authors:** Samantha H. Kannry, Sean M. O’Rourke, Suzanne J Kelson, Michael R Miller

## Abstract

Preservation of life-history and other phenotypic complexity is central to the resilience of Pacific salmon stocks. Steelhead (*Oncorhynchus mykiss*) express a diversity of life history strategies such as the propensity to migrate (anadromy/residency) and the timing and state of maturation upon return to freshwater (run-timing), providing an opportunity to study adaptive phenotypic complexity. Historically, the Eel River supported upwards of one million salmon and steelhead, but the past century has seen dramatic declines of all salmonids in the watershed. Here we investigate life history variation in Eel River steelhead by using Rapture sequencing, on thousands of individuals, to genotype the region diagnostic for run-timing (*GREB1L*) and the region strongly associated with residency/anadromy (OMY5) in the Eel River and other locations, as well as determine patterns of overall genetic differentiation. Our results provide insight into many conservation related issues. For example, we found distinct segregation between winter and summer-run steelhead correlated with flow dependent barriers in major forks of the Eel; that summer-run steelhead inhabited the upper Eel prior to construction of an impassable dam, and that both life-history and overall genetic diversity have been maintained in the resident trout population above; and no evidence of the summer-run allele in the South Fork Eel, indicating that summer run-timing cannot be expected to arise from standing genetic variation in this and other populations that lack the summer-run phenotype. The results presented in this study provide valuable information for designing future restoration and management strategies for *O. mykiss* in Northern California and beyond.

## Introduction

Anadromous salmon and steelhead are emblematic cultural symbols, particularly in the Pacific Northwest (Swezey & Heizer, 1977, Gregory et al., 2012). Indigenous communities have relied on the sustenance they provide since time immemorial. Extensive religious ceremonies arose around the seasonal returns of salmon and steelhead, aimed at maintaining their abundance in perpetuity (Swezey & Heizer, 1977). Likewise, many Euro-American fishing communities developed around the ample resources provided by the reliability of the annual returns (National Research Council, 1996). Preservation of life-history and other phenotypic complexity is central to the resilience and sustainability of Pacific salmon stocks (Hilborn et al., 2003, Carlson & Satterthwaite, Moore et al., 2010, Moore et al., 2014, Braun et al., 2016), yet genetic biodiversity loss across salmonids in the Pacific Northwest and California is estimated at 27% since European contact (Gustafson et al., 2007). Furthermore, populations in California are currently in a widespread state of collapse (Yoshiyama and Moyle, 2010, Katz et al., 2013, Moyle et al., 2017). Substantial portions of habitat loss have occurred where highly specialized adaptations have evolved (McClure et al., 2008), further compounding conservation management challenges.

Steelhead *(Oncorhynchus mykiss)* are found along the Pacific Rim from Baja California to Alaska and across to Kamchatka, Russia (McCusker et al., 2000). Natal homing has facilitated the evolution of enormous intraspecific diversity, both throughout their range and within individual watersheds (Dittman and Quinn, 1996, Katz et al., 2013). Thus, steelhead exhibit a broad spectrum of life-history patterns, which are accompanied by varying benefits and costs (Kendall et al., 2015), and lead to an array of population structuring (Papa et al., 2007). Propensity to migrate in steelhead is one readily accessible example of life-history diversity. The resident form, rainbow trout, remain in freshwater for their entire lives, but may migrate between streams and lakes or reservoirs (Holecek et al., 2012). The anadromous form rear in freshwater and embark on one or multiple ocean migrations, returning to freshwater to spawn. The migration phenotype of an *O. mykiss* is determined by a combination of environmental and genetic factors that lead to an individual condition which tends toward migration or residency (Sloat et al., 2014, Kendall et al., 2015). Across a broad geographic area, migration is associated with a large genomic region, OMY5 (Pearse et al 2014), which has two inversions in close proximity (Pearse et al., 2019). SNPs in this region of the genome are conserved together, and can be used to categorize individuals into resident, migratory, or heterozygote genotype groups, even if these genotypes do not necessarily predict migration at the individual level. This is especially true in populations that contain both migratory and resident fish such as the South Fork Eel River (Kelson et al., 2019, Kelson et al., 2020).

Another form of life-history variation in *O. mykiss*, adult “run-timing,” is largely controlled by genetics (Prince et al., 2017, Thompson et al., 2019), and presents a fascinating example of phenotypic complexity in a salmonid species. The two primary migration types in the Eel River are winter-run (also referred to as mature migrators), which enter freshwater sexually mature in the late fall through winter, spawn and return to the ocean quickly (relative to summer-run), and summer-run (also referred to as premature migrators), which enter the river sexually immature in the late spring/early summer, spend the summer and fall months maturing and awaiting the return of the rains before spawning in winter, typically high in the watershed (Jones, 1992, Busby et al., 1996). Premature migration is thought to have evolved because it opened previously unavailable temporal and/or spatial spawning habitats (Papa et al., 2007, Quinn et al., 2015). Positive selection on premature migration was likely facilitated by physical or temporal barriers that at least partially exclude mature migrators from specific habitat (Clemento, 2007, Thompson et al., 2018), but the details of this process and the specific evolutionary advantages of premature migration are imperfectly understood (Papa et al., 2007).

Premature migration is a challenging life history (Prince et al., 2017), and anthropogenic alterations to the landscape have made it more difficult. Summer-run steelhead do not feed as adults in freshwater, so they must have enough fat stored prior to migration to survive throughout harsh summer conditions and then develop gonads in preparation for spawning. Therefore, they enter freshwater with higher percent body fat than their winter-run counterparts (Smith, 1969, Hearsey and Kinzinger, 2014; Lamperth et al., 2016). Steelhead have a limited thermal tolerance (Richter and Kolmes, 2006), which is much less of a problem for winter-run, whose adult freshwater residence time is limited to the cold winter months. In contrast, summer-run adults spend the summer in cold pools (e.g., deep, thermally stratified pools) which provide refugia from high temperatures (Nielsen et al., 1994). Many of the anthropogenic changes to the landscape have direct, warming impacts on stream temperature (Poole and Berman, 2001). As a consequence of their life history strategy, summer-run (and the congeneric spring-run Chinook, *O. tshawytscha*) in Northern California have experienced severe declines in recent decades (Prince et al., 2017, Thompson et al., 2018, Waples & Lindley, 2018) due to dams, diversions and subsequent flow alteration, increasing stream temperatures, climate change, and reliance on disturbed headwaters reaches for spawning and rearing (Busby, 1996, Moyle et al., 2008, Katz et al., 2013, Arciniega et al., 2016, Quinn et al., 2016). Adaptations related to variable flows and temperature, such as premature migration by summer-run steelhead, will likely be essential to the species’ ability to persist in a changing climate (McClure et al., 2008). Increasing understanding of the ecology and historical evolutionary advantages experienced by summer-run steelhead will help promote their persistence.

The Eel River in Northwestern California is currently the southernmost river with summer-run steelhead in North America (Jones, 1992). The Eel River is also the most erosive non-glacial river in North America, creating a dramatically unstable landscape (Power et al., 2015, Roering et al., 2015) of boulder roughs (accumulations of house sized boulders in steep constrained channels, ranging 0.25-5.00 km long, frequently at the foot of massive landslides and shifting annually), with varying passage potential for migrating salmonids. Historically, the Eel River supported upwards of one million salmon and steelhead and was home to one of the largest recreational and commercial fisheries on the West Coast of the United States (Yoshiyama and Moyle, 2010). Presently, three of the four salmonid species in the Eel River, steelhead, Chinook, and coho salmon *O. kisutch*, are listed as threatened under the USA Endangered Species Act (ESA). Current population estimates in the Eel River represent a 99% decline from historical figures (Jones, 1992, Yoshiyama and Moyle, 2010). The primary causes of decline in the Eel River are the same as those across the range of salmonids (Yoshiyama and Moyle, 2010), dams and diversions, overfishing, poor logging practices, human development and habitat loss, grazing impacts, hatcheries and climate change (Katz et al., 2013). Eel River populations have recently been threatened again by the marijuana “green rush” and its associated land conversion, summertime water extraction and warming temperatures in natal streams (Carah et al., 2015).

Genetic analysis of the run-timing phenotypes in the Middle Fork Eel River (and other rivers) found that premature and mature migrating steelhead are more closely related to each other than they are to populations with the same phenotype in other basins (Nielsen and Fountain, 1999, Clemento, 2007, Arciniega et al., 2016, Prince et al., 2017). These results were interpreted to suggest that the summer-run phenotype evolved independently in different rivers and, if extirpated, could rapidly re-evolve from winter-run individuals (Thorgaard, 1983, Waples et al., 2004, Pearse et al. 2020). However, recent research suggests that run-timing in steelhead (and Chinook) is controlled by a single locus (i.e., the *GREB1L* region), and the summer-run allele evolved once and then spread throughout their range (Prince et al., 2017). Thus, the summer-run phenotype cannot be expected to rapidly re-evolve from winter-run populations (Prince et al., 2017). Furthermore, as the premature migration phenotype is extirpated, the allele will not be maintained by mature migrating populations because heterozygotes have an intermediate migration phenotype (Prince et al., 2017; Thompson et al., 2018).

Here we examine a number of issues related to steelhead conservation in the Eel Basin. Our primary goal is to address the scarcity of information on the ecology and distribution of summer-run steelhead. More specifically, we explore population structure and demography across the Eel basin, the spawning and rearing distribution of the two adult run-timings (as well as heterozygotes), the distribution of resident and anadromous genotypes to inform the extent to which upstream migration may be inhibited by particular geographical features, and the state of genetic variation and diversity in a population of resident trout isolated above an impassable dam for nearly a century. To this end, we collected tissue samples from *O. mykiss*, primarily juveniles, in the Van Duzen (n=478), Middle Fork (n=183) and upper mainstem Eel (n=173) Rivers (Figure 1 & Table 1, S1) from June 2016-October 2018 and analyzed them using Rapture sequencing (Ali et al. 2016), targeting both loci spread across the genome and those linked to life-history variation (i.e., *GREB1L* and OMY5). Furthermore, we also analyzed Rapture data from the South Fork Eel (n=2089, Table S2) collected for a previous study (Kelson et al., 2019, Kelson et al., 2020) to investigate the extent to which the summer-run allele is maintained as standing variation in winter-run populations.

**Table 1.**
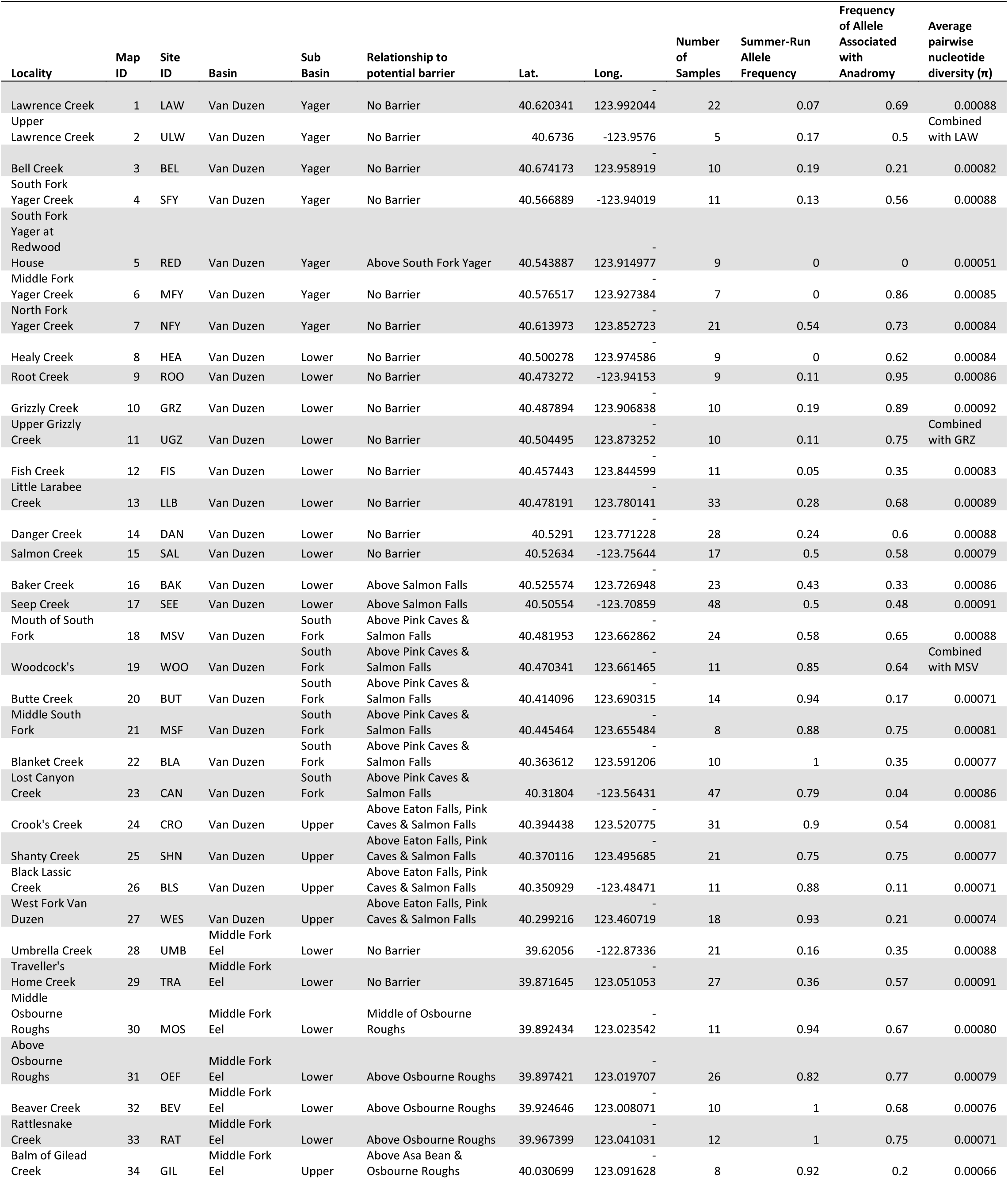
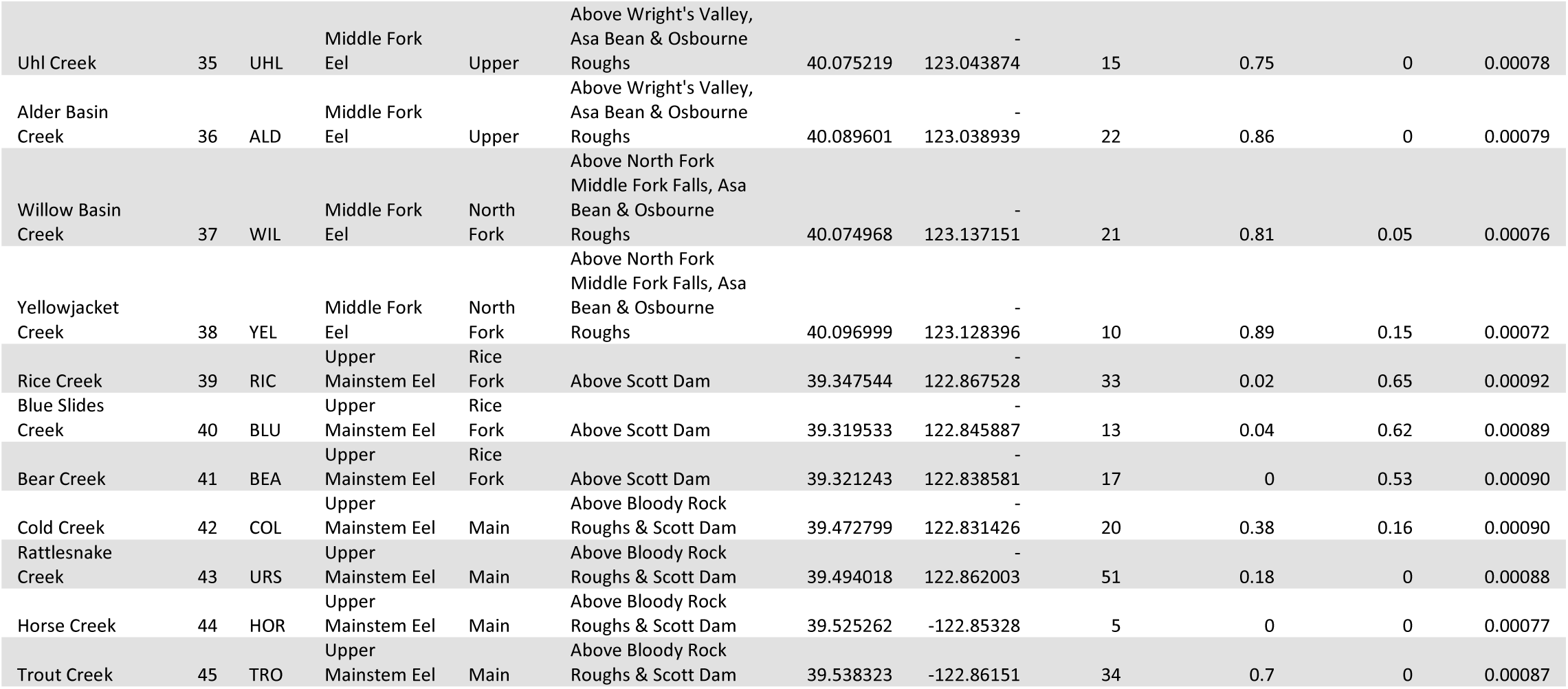
Sampling location information

**Figure 1.**
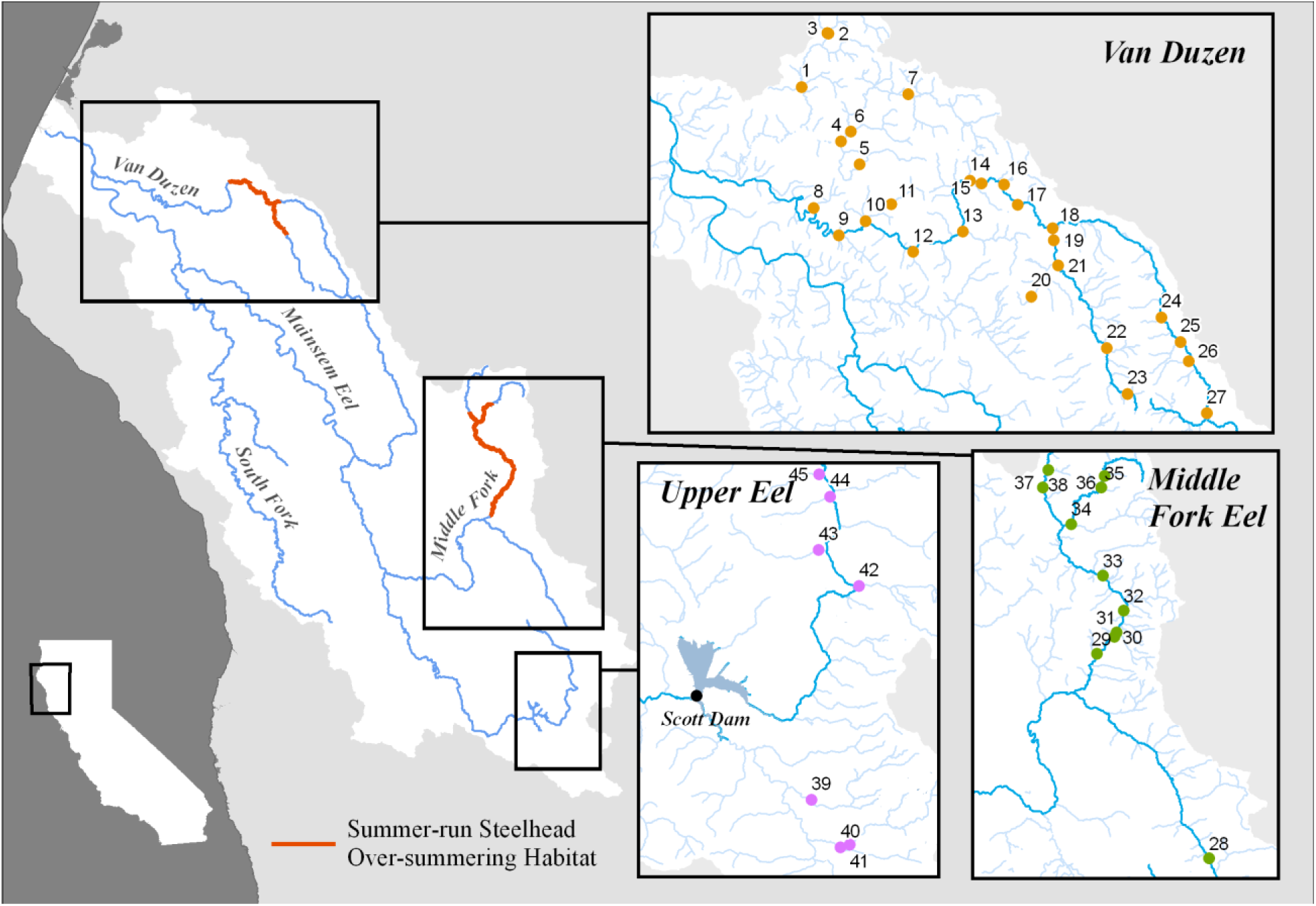
Map of Eel River watershed with sampling locations in the three basins indicated as dots and numbered (numbers referenced in Table 1). Current summer-run steelhead summertime holding habitat highlighted in orange.

## Results

### Population structure analysis reveals two discrete summer-run populations in the Eel River

Winter and summer-run steelhead in the Eel River are part of the Northern California Distinct Population Segment, which extends from the Gualala River in Sonoma County to Redwood Creek in Humboldt County and inland to the headwaters of the Eel River in Lake County (NMFS, 2016). Currently, there are two locations in the Eel Basin where summer-run individuals are routinely observed: the Van Duzen and Middle Fork Eel Rivers, two of the major forks of the Eel River (Figure 1). Together, they presently constitute the southernmost expression of this vulnerable phenotype in North America (Nielsen and Fountain, 1999). Since 1966, California Department of Fish and Wildlife (CDFW) has monitored adult summer steelhead abundance in the Middle Fork Eel by conducting an annual direct observation census of all summer holding habitat. The result of this census is an index of anadromous adult abundance with a mean size of 770 individuals per year (Harris, 2019). A comparable census was implemented in the Van Duzen River five times between 1979 and 2011, yielding a mean size of 25 individuals per year. However, in 2011, a consistent annual census began and has resulted in a population count ranging from 51-255, with a mean of 148 individuals per year (Harris, 2019). These surveys suggest both locations have experienced a dramatic reduction in population size from historical abundance, and with it the potential loss of genetic variation (McClure et al., 2008) and jeopardized long-term viability (Yoshiyama and Moyle, 2010). Previous genetic analyses of summer-run steelhead in the Eel have not included the Van Duzen population (Nielsen and Fountain, 1999, Clemento, 2007), and the small observed population size of Van Duzen adult summer steelhead has led to speculation that the spawners documented in the sub-basin are strays from the Middle Fork Eel population as opposed to an independent population.

To investigate population structure in the Eel River with an emphasis on the origin of Van Duzen summer steelhead, we performed a principal component analysis (PCA) on a subset of the samples that provided an even representation of the Van Duzen, Middle Fork Eel, and Upper Eel Basins (n=118 for each basin, Table S1, Figure 2) (Materials and Methods). This and other downstream analyses were done with Young of the Year (YOY) with a few notable exceptions (Materials and Methods). Samples clustered into three main groups corresponding to geography (as opposed to migration phenotype), with samples collected from the lower reaches of the basins being well-mixed and increased upstream differentiation for each basin. In other words, summer-run fish cluster with winter-run fish in both the Van Duzen and Middle Fork Eel Rivers, as opposed to summer-run fish from the Van Duzen clustering with Middle Fork Eel fish, which would be expected if they were strays. We conclude that summer-run steelhead in the Van Duzen River represent an independent population, not a population maintained by dispersal from the Middle Fork Eel River.

**Figure 2.**
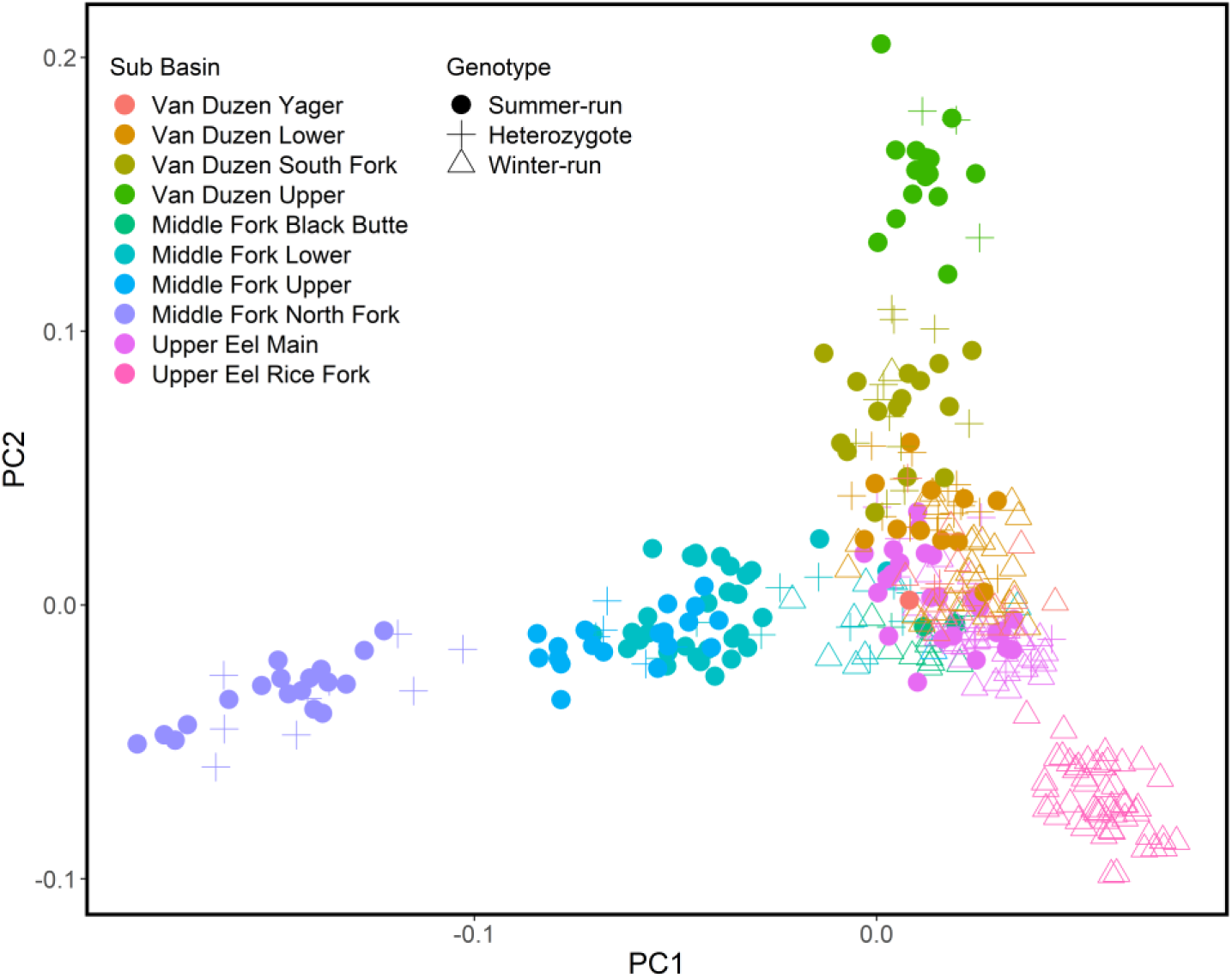
Principal components analysis of individual genotype data from a subsample of YOY individuals collected across the three basins (n=118 for each basin). PCA was run on all sequenced markers, excluding chromosome 5. Color indicates sub-basin where individual was sampled, and shape indicates run-timing of individual based on GREB1L locus genotype.

### Eaton Falls is not a total barrier to upstream migration

Geographic barriers to fish passage have been observed to substantially alter the dynamics of local populations of steelhead and result in genetic divergence of neighboring populations (Pearse et al., 2009, Limborg et al., 2011, Martinez et al., 2011). The geographical feature termed “Eaton Falls” on the Van Duzen River is at the upstream end of a half-kilometer stretch of steep roughs that gains 90 meters in elevation, culminating with an eight-meter bedrock wall at Eaton Falls. Along the left bank of the channel is a jumble of house sized boulders that is partially inundated at high flows. Above the falls, there are no mainstem barriers to migration for the subsequent 50 km, until the river becomes too high gradient for anadromous fish passage (Nethery, 1973). Although the section of river above the falls is managed as non-anadromous waters by state and federal agencies (Nethery, 1973), a recent study conducted isotope analysis on 11 otoliths collected from juvenile *O. mykiss* above the falls and found one individual with anadromous maternal origin (B. Harvey, unpublished data). Furthermore, in mid-August of 2018 two authors of this manuscript observed a 58 cm adult summer-run steelhead 1 km above Eaton Falls. These observations suggest that Eaton Falls may not be a total barrier to migration of adult steelhead. Another geographical feature considered to be a complete barrier to anadromy is found on the South Fork of Yager Creek, major tributary to the Van Duzen River. These are the two major features on main forks of the Van Duzen above which anadromous individuals are not routinely observed, here termed “potential complete barriers” (PCBs) (Table S3). Therefore, they serve as valuable locations to compare in terms of passage potential for steelhead.

The resident and anadromous life histories in steelhead have been shown to be strongly associated with the OMY5 region (i.e., a large region on chromosome 5) in California and in the Eel River, specifically (Pearse et al., 2014, Kelson et al., 2019; Pearse et al. 2019). A number of studies have observed higher frequencies of resident genotype individuals above natural and man-made barriers than below them (Pearse et al., 2014, Abadia-Cardoso et al., 2016, Phillis et al., 2016, Leitwein et al., 2017). The anadromous variant in the OMY5 region has been shown to be lost in some populations above barriers, unless the populations have access to a reservoir or lake and behave in an adfluvial way (Holecek et al., 2012; Leitwein et al., 2016 and Arostegui et al., 2019). In addition, anadromy may be selected against when food resources are plentiful and the fitness cost to migration outweighs the benefits (McClure et al., 2008). Therefore, the genotype frequencies at the OMY5 region can serve as a proxy for the potential for anadromy at a given location, especially if there is no reservoir or lake to maintain adfluvial migrations, which is the case above Eaton Falls.

To investigate the degree to which Eaton Falls inhibits fish passage, we genotyped the Van Duzen samples at the OMY5 region (n=352) and created a genotype frequency map (Table 1 & Figure 3; Materials and Methods). We did not detect a substantial shift in OMY5 allele frequency between samples collected above and below Eaton Falls. For example, the frequencies of the allele associated with anadromy observed at the four locations sampled above Eaton Falls, from downstream to upstream, Crooks Creek (0.54), Shanty Creek (0.75), Black Lassic Creek (0.11) and West Fork Van Duzen (0.21), follow a similar pattern to that observed in the South Fork Van Duzen, which is not above a barrier to anadromy and routinely used by anadromous adults for spawning (Table 1 & Figure 3). Another useful comparison are the three sites below Eaton Falls, from downstream to upstream, which have the following frequencies for the allele associated with anadromy, Baker Creek (0.33), Seep Creek (0.48) and Mouth of South Fork Van Duzen (0.65) (Table 1). Furthermore, we found that above the potential barrier to anadromy on the South Fork of Yager Creek, the population is fixed for the allele associated with residency, confirming that the OMY5 locus follows similar patterns in the Van Duzen as other locations (Pearse et al., 2014, Abadia-Cardoso et al., 2016, Leitwein et al., 2017). We conclude that unlike the South Fork Yager feature, Eaton Falls does not appear to be a complete barrier, as the frequency of the allele associated with anadromy above Eaton Falls is higher than would be expected if the falls were impassable to steelhead.

**Figure 3.**
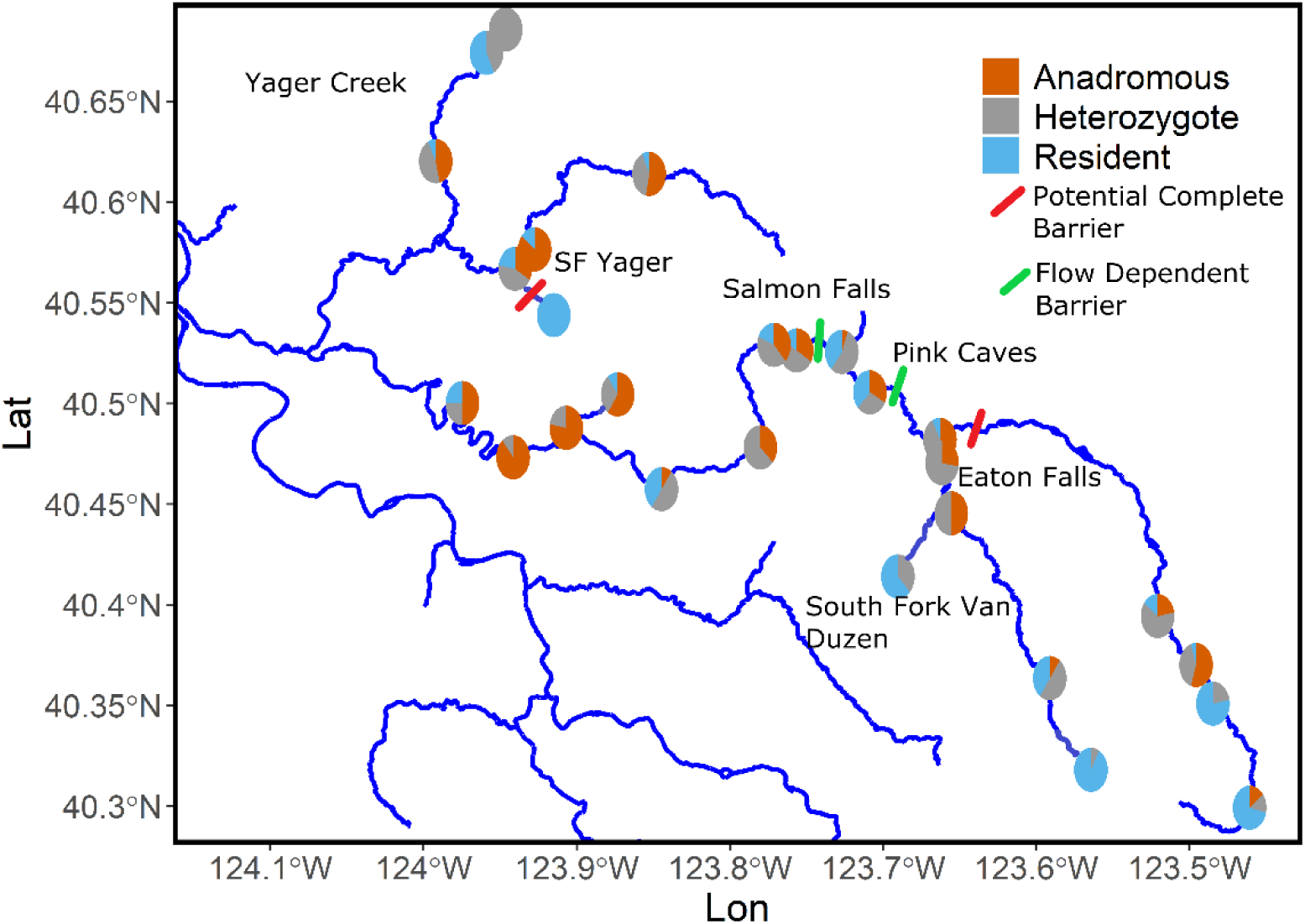
Map of the Van Duzen River depicting YOY OMY5 genotype distribution. Circles represent sampling location and colors indicate genotypes frequency at the location. Barriers are indicated by bars and color of the bar represents the type of barrier. All barriers are natural barriers.

To investigate the extent to which Eaton Falls reduces gene flow, we estimated pairwise F_ST_ for all site combinations in the Van Duzen River, and compared the pairwise estimates that do not cross a barrier to those that cross Eaton Falls and the South Fork Yager barrier (Figure 4). The mean pairwise F_ST_ values were 0.059 (range, 0.018 to 0.149; n=177) for the site combinations that do not cross a barrier, 0.094 (range, 0.052 to 0.160; n=76) for the site combinations that cross Eaton Falls, and 0.219 (range, 0.173 to 0.268; n=23) for the site combinations that cross the South Fork Yager barrier (Table S4). Permutation testing revealed that the increase in pairwise F_ST_ values for both sets of site combinations (those crossing Eaton Falls and those crossing the South Yager Barrier) were significantly elevated relative to the site combinations that do not cross a barrier (p<0.05). Taken together, our results suggest that Eaton Falls does not represent a total barrier to upstream migration and anadromous steelhead likely pass it with at least some regularity.

**Figure 4.**
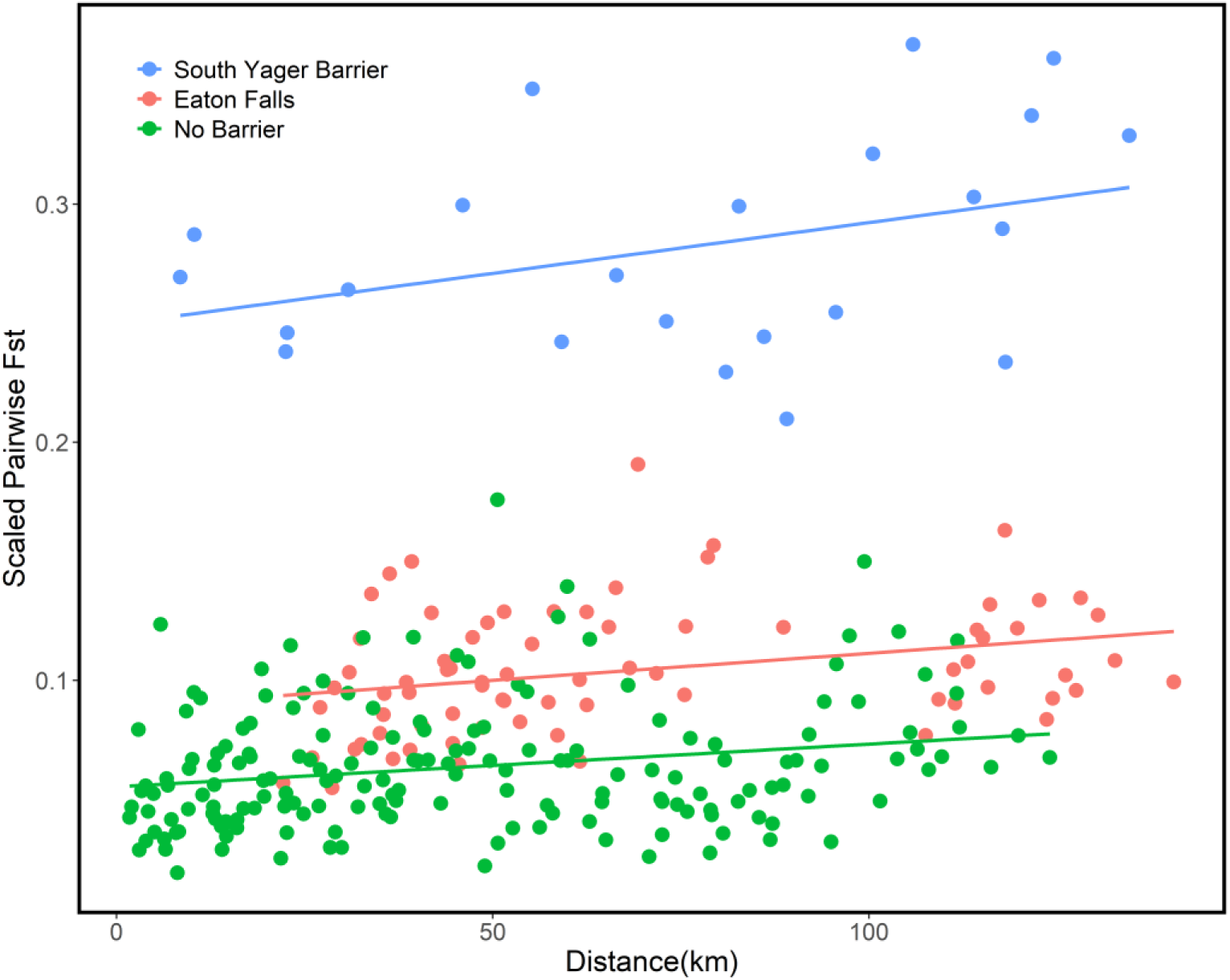
Isolation by distance plot of Van Duzen pairwise combinations. Each point represents a pairwise comparison of sites on the Van Duzen. Color indicates the relationship of the two sites being compared to the two potential complete barrier being investigated (e.g., Eaton Falls means Eaton Falls is between the sites).

### Distinct juvenile distribution between runs correlated with flow dependent barriers

Summer-run steelhead may be able to navigate through barriers frequently impassable during winter-run migration times, due to lower (but not excessively low) and consistent late spring and early summer flows. However, this benefit is tempered by fitness trade-offs such as increased adult mortality in extended freshwater residence (Hess et al., 2016). While both spatial and/or temporal differences in migration and spawning have been proposed to segregate premature and mature migrating populations of steelhead and Chinook (Nielsen and Fountain, 1999, High et al., 2006, Papa et al., 2007, Hess et al., 2016, Thompson et al., 2018), there has been little research into the influence physical barriers have on spawning and rearing distributions of the two run-timings. Thus, it remains unclear if strong spatial or temporal barriers exist between spawning summer-run and winter-run fish on the Van Duzen and Middle Fork Eel Rivers. The over-summering habitat of steelhead in the Eel River is characterized by long steep stretches of roughs, interspersed with cobble and gravel bars, which can prevent passage to summer steelhead late in their migration period (e.g., late June or July) if flows become too low, isolating fish in pools at the downstream ends of the roughs. Examples of this are found at the Asa Bean Roughs on the Middle Fork Eel and Salmon Falls on the Van Duzen, both of which are regularly observed to strand fish in the late summer and early fall (Harris, 2004), here termed “flow dependent barriers” (FDBs) (Table S3). The other barriers examined in this study present navigational obstacles to migrating salmonids, but are either near complete barriers at all flows (Wright’s Valley and North Fork Middle Fork, both on the Middle Fork Eel and PCBs) or have not been observed to strand fish in the summer (Osbourne Roughs on the Middle Fork Eel and the Pink Caves on the Van Duzen, both FDBs) (Table S3).

To examine the distribution of summer and winter-run spawning and rearing in the Van Duzen and Middle Fork Eel Rivers and determine the extent to which different types of barriers impact passage and the spatial delineation of the two runs, we collected juvenile samples from multiple locations in both rivers, upstream and downstream of FDBs and PCBs. We analyzed them with markers in the *GREB1L* region that appear diagnostic for run-timing (Prince et al. 2017; Materials and Methods) and generated genotype frequency maps. We observed a dramatic partition in the frequency of the run-timing genotypes as we travel from downstream to upstream past FDBs in the Van Duzen River (Figure 5, Table S5). Below the Salmon Falls barrier, we see primarily individuals with the winter-run genotype. As we move into the summer-run holding habitat, known as the Lost Duzen region, we see higher frequencies of heterozygotes and similar frequencies of both winter and summer-run genotypes. Above the Pink Caves and Eaton Falls, we found primarily individuals with summer-run genotypes, with some heterozygotes and a very low frequency of winter-run genotypes. We found an analogous pattern in the Middle Fork Eel (Figure 6, Table S5), although at a less fine spatial scale due to sampling limitations (Materials and Methods), with Osbourne Roughs being the primary barrier between the two runs. We conclude that summer and winter-run steelhead use distinct spawning and rearing habitat in both the Van Duzen and Middle Fork Eel Rivers, with winter-run fish largely excluded from the habitat above FDBs.

**Figure 5.**
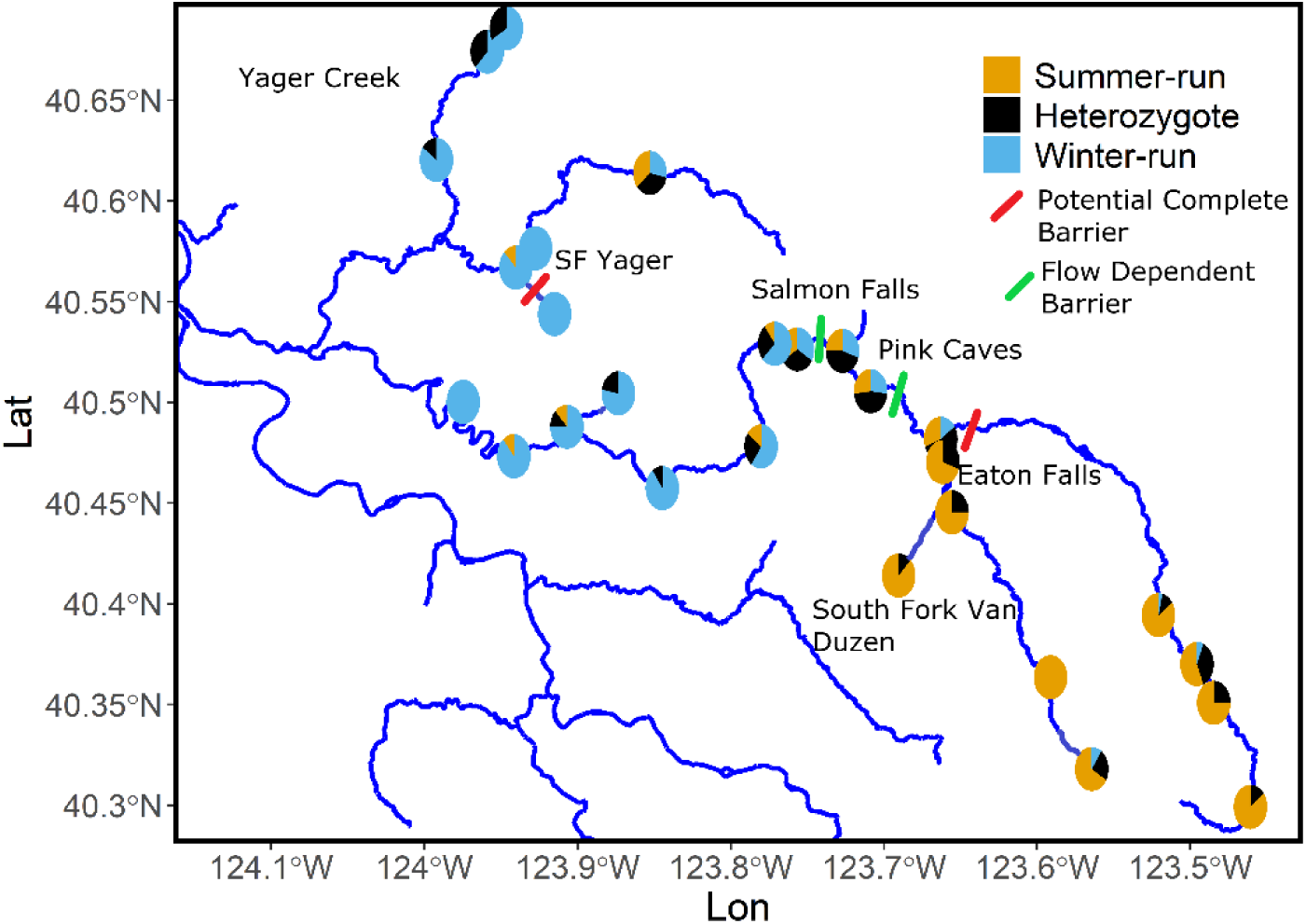
Map of Van Duzen River YOY GREB1L genotype distribution. Circles represent sampling location and colors indicate genotypes frequency at the location. Barriers are indicated by bars and color of the bar represents the type of barrier. All barriers are natural barriers.

**Figure 6.**
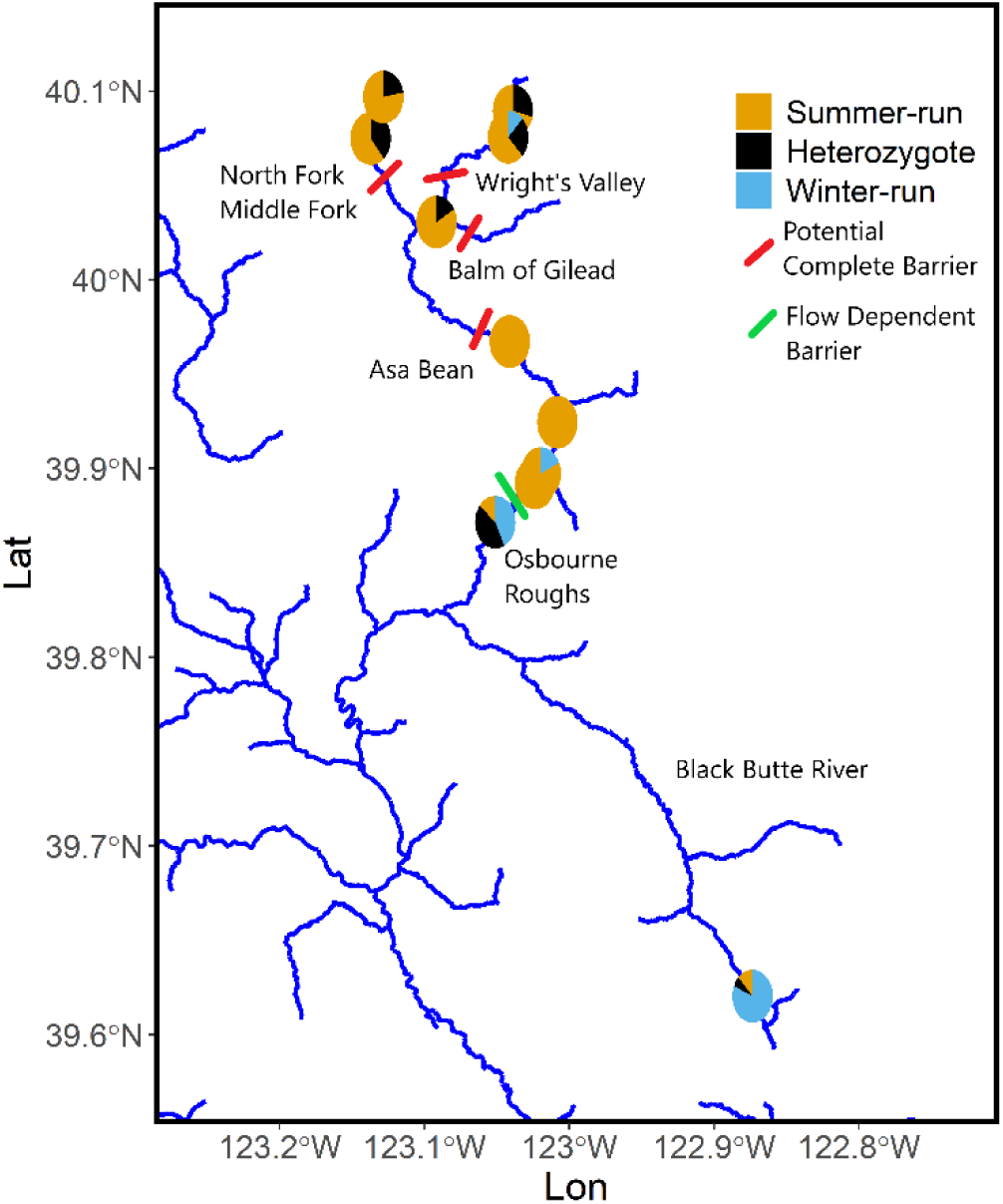
Map of Middle Fork Eel River YOY GREB1L genotype distribution. Circles represent sampling location and colors indicate genotypes frequency at the location. Barriers are indicated by bars and color of the bar represents the type of barrier. All barriers are natural barriers.

### Resident trout population above Scott Dam equipped for re-establishing an anadromous population

Salmonids are presently excluded from almost 45% of their historic range in the lower 48 states due to severe habitat alterations and anthropogenic barriers (McClure et al., 2008). The two dams on the Eel River, Cape Horn and Scott Dams, impound 890 km^2^ of the upper mainstem Eel River. Cape Horn Dam has fish passage and was built first in 1907, however the dam’s reservoir quickly filled with sediment leading to construction of the upstream Scott Dam. Scott Dam was built with no fish passage and has blocked steelhead access to 179-291 km of rearing and 318-463 km of spawning habitat (Cooper, 2017). Although constructed as a hydropower facility, the project’s current primary function is diverting water through a tunnel, south into the Russian River watershed for agricultural and residential use (Power, 2015). The year 2022 will mark the second 50-year re-licensing cycle under the Federal Energy Regulatory Commission (FERC). Due to financial, liability, fish passage and feasibility issues, there is potential for removal of Scott Dam, restoring anadromous fish access to the upper portion of the watershed after 100 years. Although resident rainbow trout remain above the dam, small and disconnected populations above dams can experience decreases in genetic diversity (Clemento et al., 2009), loss of the anadromous life history variant (Pearse et al., 2014), or, conversely, they can maintain the migratory life-history in an ad-fluvial form (Holecek et al., 2012; Leitwein et al., 2016 and Arostegui et al., 2019). In addition, there is a lack of clear evidence as to whether or not there was a population of summer-run steelhead in the area that is now blocked from migrating salmonids by Scott Dam.

To investigate the genetic attributes of resident rainbow trout above Scott Dam, we collected 173 tissue samples from the main and Rice forks of the Eel River above the dam and genotyped them at *GREB1L*. Interestingly, we found individuals carrying the summer-run alleles in three tributaries of the main fork, Cold Creek (COL, homozygous summer-run=0.25, heterozygote=0.25), Rattlesnake Creek (URS, homozygous summer-run=0.17, heterozygote=0.02) and Trout Creek (TRO, homozygous summer-run=0.60, heterozygote=0.20), (Figure 7 & Table 1). We also found two heterozygotes in the 57 samples we collected from three tributaries of the Rice Fork. These results match what we would expect to see both ecologically and geographically, as the upper main fork and its tributaries originate from higher elevation spring-fed sources (Murphy and Rodriguez, 1993), remain colder throughout the summer (Crawford, 2017), and have steeper gradients and deeper pools that could have provided over-summering habitat for adult summer steelhead (Murphy and Rodriguez, 1993). In contrast, the Rice Fork is low gradient, flows intermittently in the summer, and regularly has temperatures in the lethal range for steelhead (Murphy et al., 1993, Crawford, 2017). We conclude that summer-run steelhead inhabited the upper Eel, prior to construction of Scott Dam in 1922 and subsequent obstruction of upstream passage, and the summer-run alleles have up to now persisted in the resident population.

**Figure 7.**
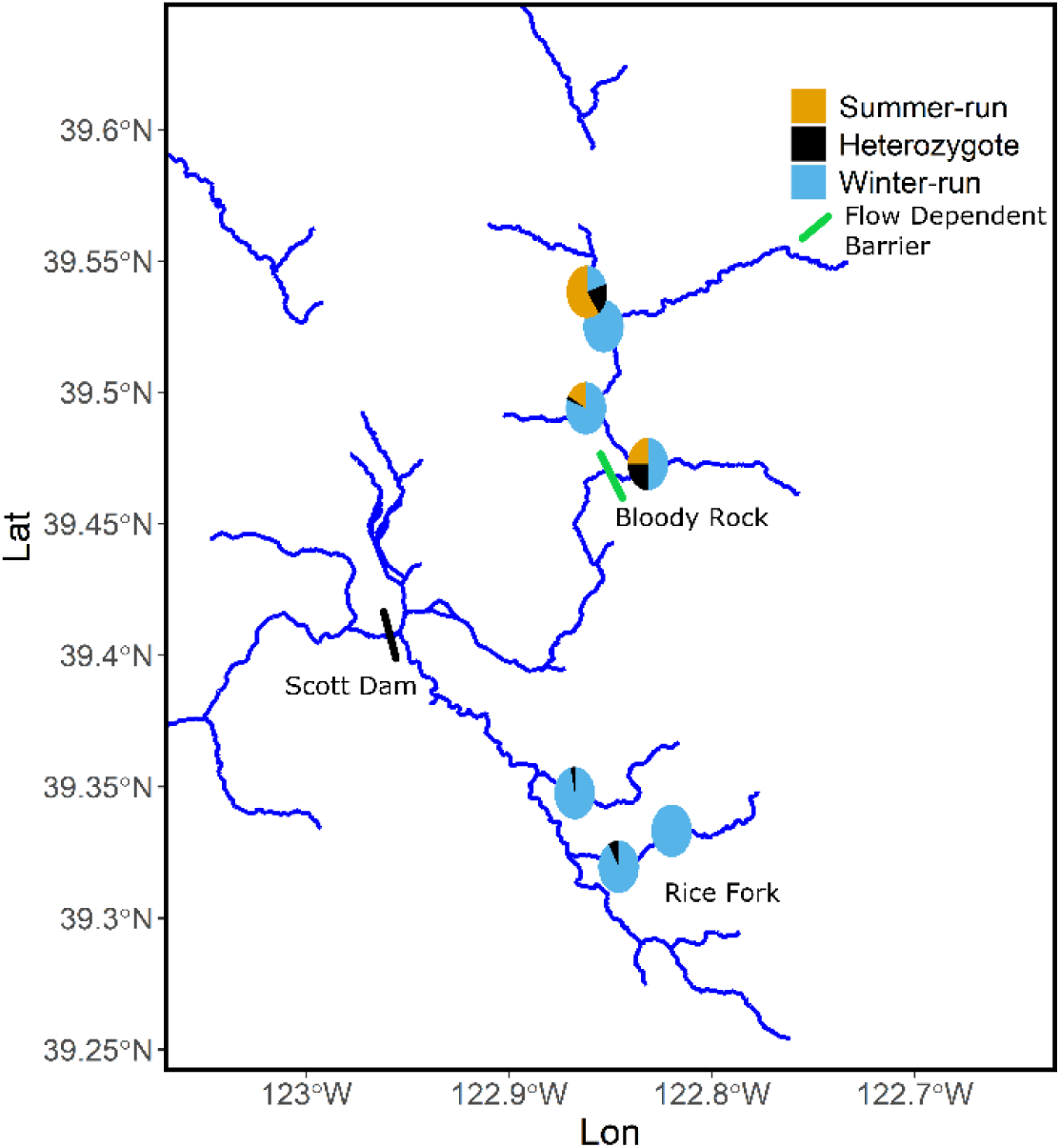
Map of Upper Eel YOY GREB1L genotype distribution. Circles represent sampling location and colors indicate genotypes frequency at the location. Barriers are indicated by bars and color of the bar represents the type of barrier. Bloody Rock is a natural barrier and Scott Dam is a man-made barrier.

To investigate the genetic variation with respect to migratory potential of resident trout above Scott Dam, we genotyped the individuals from the upper Eel at the OMY5 region. We called genotypes (n=156) as above and created a map of genotype frequencies (Figure 8). We found primarily individuals with the genotype associated with residency on the main fork above the one obstruction in that region, Bloody Rock Roughs (FDB) (Table S3) (Cooper, 2017), and primarily heterozygous individuals and individuals with the genotype associated with anadromy in the Rice Fork. While Bloody Rock does not inhibit steelhead migration, resident trout may not attain sufficient size to migrate upstream, through this type of steep roughs, if they expressed an adfluvial life-history (Holecek et al., 2012). We conclude that genetic variation with respect to migratory potential has, up to the present, been maintained in a subset of the resident trout above the dam.

**Figure 8.**
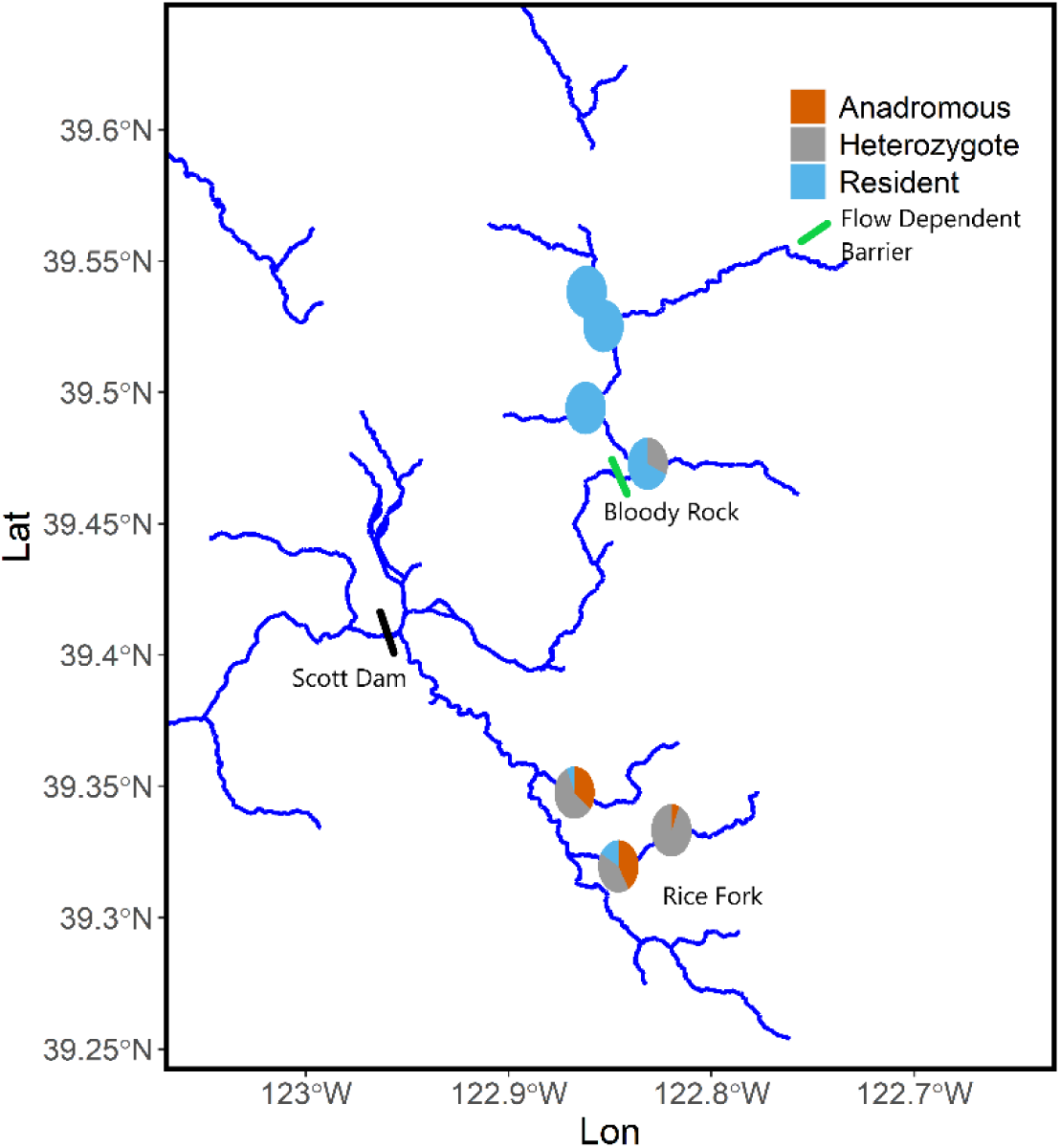
Map of Upper Eel YOY OMY5 genotype distribution. Circles represent sampling location and colors indicate genotypes frequency at the location. Barriers are indicated by bars and color of the bar represents the type of barrier. Bloody Rock is a natural barrier and Scott Dam is a man made barrier.

To examine overall genetic diversity in rainbow trout above Scott Dam, in relation to other regions of the Eel River, we calculated the average number of pairwise differences (π) (Tajima, 1983), a standard measure of genetic diversity, for all sampling locations. The mean π for each of the Middle Fork, Upper Eel and Van Duzen was 0.00078, 0.00087 and 0.00082, respectively (Figure 9, Table 1). The mean π for the upper Eel above Scott Dam was significantly higher than the Middle Fork (p<0.05), but not significantly higher than the values we observed in the Van Duzen (p>0.1). We conclude that the resident trout population above the dam has maintained overall genetic diversity. Taken together, our results suggest that resident trout populations above the dam would likely be a suitable source population for recolonization of the upper basin by anadromous individuals if the dam were to be removed.

**Figure 9.**
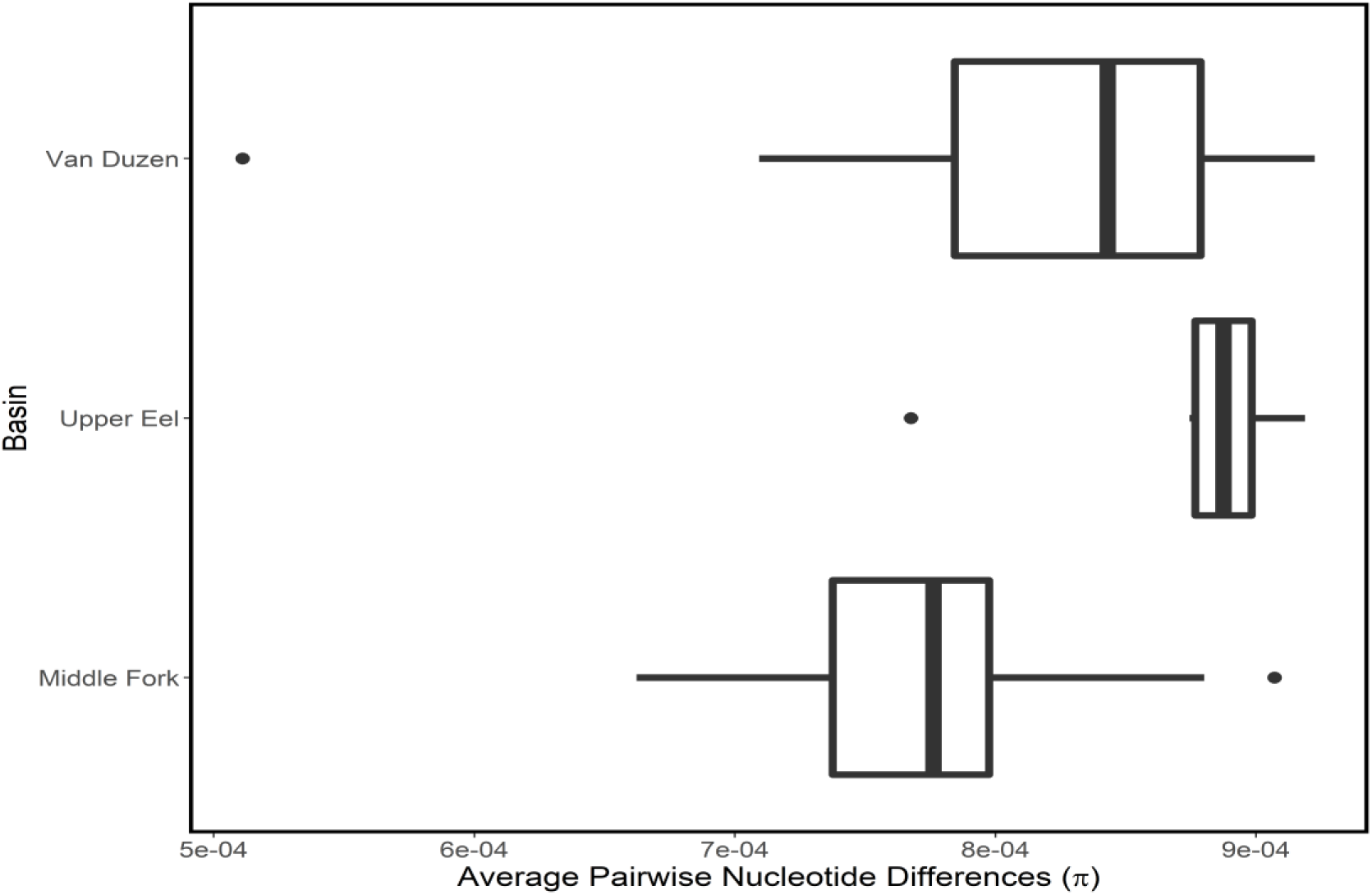
Box and whisker plot representing overall genetic diversity by basin (π).

### Summer-run alleles are not detected in the South Fork Eel River

Historical documentation suggests that summer-run steelhead inhabited the upper South Fork Eel, prior to a major flood in 1964 (Jones, 1992). The flood resulted in substantial habitat degradation and further reduced most of the already severely declining native fish stocks in the Eel River (Yoshiyama and Moyle, 2010). The summer-run fish were purported to hold in deep pools in a stretch of river surrounding what is now the UC Angelo Reserve. The evidence of a population of summer-run steelhead in the South Fork is primarily based on anecdotal reports from locals and hatchery staff in the 1950’s and 60’s. Summertime snorkel surveys in the 1970’s did not observe adult steelhead (Jones, 1992). The proposed holding habitat is composed of a bedrock-lined gorge, but does not contain the steep roughs (Elwell and Fisk, 1959) that function as spatial barriers between winter and summer-run steelhead in the other Eel River sub-basins (see above). Furthermore, the headwaters of the South Fork Eel are relatively low in elevation (Trush et al., 1985) and receive negligible snow-pack, resulting in less consistent spring flows than the Middle Fork Eel and Van Duzen Rivers and less late spring/early summer upstream migration opportunities for summer-run steelhead.

To address the question of whether summer-run alleles are currently present in the South Fork Eel River, we genotyped 2090 individuals at the *GREB1L* locus. The samples were collected for a previous study (Kelson et al., 2019, Kelson et al., 2020) from the Elder, Fox and mainstem South Fork watersheds between 2014-2017, primarily in the Angelo Reserve area. Of the 2089 samples, 1593 had sufficient reads to make a genotype call (Materials and Methods). Strikingly, all 1593 individuals were homozygous for the winter-run genotype (Table 2, S2). Elder and Fox Creeks are in the upper portion of the watershed, where the majority of the anecdotal evidence of summer-run fish originates. We conclude that the summer-run allele is not maintained as standing variation in winter-run steelhead populations.

**Table 2.** Genotypes of individuals sampled from the South Fork of the Eel River.

| Year | Homozygous winter | Heterozygous | Homozygous summer |
| --- | --- | --- | --- |
| 2014 | 26 | 0 | 0 |
| 2015 | 550 | 0 | 0 |
| 2016 | 595 | 0 | 0 |
| 2017 | 422 | 0 | 0 |
| Total | 1593 | 0 | 0 |

## Discussion

Species are being lost at an unprecedented rate (Barnosky, et al., 2011, Ceballos, 2015). As difficult as it is to catalog and quantify the species going extinct, the local adaptations and intra-specific diversity that is vanishing may have even more severe ramifications (Mimura et al., 2017, Thompson et al., 2019). For many years, it was assumed that the Eel River had only one remaining independent summer-run steelhead population on the Middle Fork, so that is where emphasis in monitoring and research was placed. However, annual dive surveys since 2011 established that the Van Duzen harbored more individuals than previously thought. Our results clarify that the Van Duzen is an independent population as opposed to being strays from the Middle Fork Eel population. Since the extirpation of an independent summer-run population is much more consequential than the extirpation of strays, the Van Duzen summer-run populations must be considered a critical component of, and recovery source for the Northern California steelhead Distinct Population Segment. Furthermore, since its numbers are quite low and it is not likely to re-evolve from winter-run steelhead (Prince et al., 2017; see below), the Van Duzen population is confronting a substantial risk of extirpation.

We further confirmed that steelhead are capable of surprising feats, such as migrating through alleged total barriers to migration like Eaton Falls on the Van Duzen River. We found both relatively high frequencies of the genotype associated with anadromy (OMY5, Pearse et al., 2014, Kelson et al., 2019), and a relatively minor increase in F_ST_ values in pairwise site combinations crossing the falls compared to those that did not. These findings, in conjunction with previous observations of summer-run steelhead above Eaton Falls, provides further support for the assertion that Eaton Falls is not a complete migration barrier. Since the Van Duzen River above Eaton Falls is managed as non-anadromous waters, fishing regulations are relaxed, as are restrictions on permitting for development in the floodplain. These results suggest that even accepted total barriers can be passable with some frequency. Considering the instability of the slopes adjacent to the Van Duzen River (Roering et al., 2015), Eaton Falls may fluctuate between degrees of pass-ability over relatively short time scales. Such has been observed to be the case on the Middle Fork Eel River at Asa Bean Roughs. Prior to 2011, the roughs were passable annually, earlier in the season and would only prevent passage to the late-season migrants. During the winter of 2011, some combination of precipitation and/or other geological forces caused a shift in the placement of a few key boulders at the lower end of the roughs, resulting in a complete barrier to migration. Since then no adult anadromous steelhead have been observed in the annual dive surveys above the roughs and juvenile densities in the previously anadromous-accessible stretch above Asa Bean have decreased dramatically. An index site approximately 6 km upstream of the roughs, surveyed since 1980 to estimate juvenile density has had counts of zero since 2014, down from an average density of 0.81 fish/m² (Harris, 2004, Harris, 2019). Summer-run steelhead are no longer able to access approximately 15 km of mainstem and additional tributary habitat for rearing and spawning, a considerable loss with the already severely depressed population size.

Consistent receding spring flows typically provide summer-run fish passage to upper portions of the watersheds (Bjorkstedt et al., 2005). In the Van Duzen, Middle Fork, and upper Eel watersheds, snowmelt contributes to the spring flows but is likely to decrease in the coming decades due to climate change (Miller et al., 2003) because the watersheds are located relatively far south and have fairly low maximum watershed elevations, 1803m, 2467m and 2151m, respectively. Reduced spring flows from decreased snowpack and/or other anthropogenic disturbances combined with the confounding issue of inconsistent upstream passage through the roughs has the potential to reduce the historical advantage of premature migration (Hess et al., 2016; Thompson et al., 2019). This likely has already occurred, leading to the discordant decreases in population sizes of summer-run steelhead, relative to winter-run. A primary goal of this study was to investigate the distribution of run-timing genotypes in the Eel River. Although summer-run steelhead may have historically existed in the Sacramento-San Joaquin drainages (Swezey and Heizer, 1977), the Eel River is currently the southernmost river with a population of summer-run in North America (Busby et al., 1996). While physical barriers have been hypothesized to be a major driver in maintaining the two run-timings within a single watershed (Shapovolov and Taft, 1954, Waples et al., 2004, Quinn et al., 2015, Hess et al., 2016), we wanted to directly investigate the role barriers play in delineating spawning and rearing distribution of the two runs. Our results demonstrate that in the Eel River, summer and winter-run steelhead exhibit strong spatial segregation. Summer-run fish are primarily utilizing the upper portions of the basins, above flow dependent barriers. We did find a few heterozygotes and homozygous winter-run fish above these barriers. As the barriers shift from year to year and passable flows occur in different months, the occasional winter-run or heterozygous adult is able to reach the spawning grounds above. While this partial sympatry may have always been the case, it is likely that the severely depressed state of the summer-run populations, relative to the winter-run, has resulted in a higher frequency of heterozygotes than would have occurred historically (Ford et al., 2020) The ability to consistently pass upstream of these barriers is necessary to maintain the evolutionary advantage garnered by being a summer-run steelhead. When barriers shift as in the case of Asa Bean Roughs or anthropogenic activities reduce or eliminate the spatial segregation between the premature and mature migration phenotypes, as in the case of Lost Creek Dam on the Rogue River (Thompson et al., 2019), the advantage of being a premature migrator is reduced and will lead to a corresponding reduction or elimination of the premature migration phenotype.

The summer-run life history requires particular habitat and flow characteristics to be successful, likely resulting in minimal straying from natal sites to ensure reproductive success (Bjorkstedt et al., 2005). Our results show that the area above Osbourne Roughs on the Middle Fork Eel and above the Pink Caves on the Van Duzen are critical habitat for summer-run steelhead. The South Fork Van Duzen is the only tributary in between Salmon Falls and Eaton Falls with significant steelhead spawning habitat. The upper portion of the South Fork Van Duzen watershed is surrounded by the Mount Lassic Wilderness and the Six Rivers National Forest. One summer steelhead was observed in September 2018 (S. Kannry, personal observation), prior to the onset of fall rains and the subsequent flow increases, in the vicinity of Lost Canyon Creek, approximately 16 km above the reported South Fork Van Duzen holding habitat (Jones, 1992). The lower portion of the South Fork Van Duzen is a conglomeration of private holdings ranging from 20-acre parcels to multiple thousand-acre ranches. The major tributary to the South Fork, Butte Creek, has similarly framed land ownership with the upper portion held by the Bureau of Land Management. The South Fork Van Duzen appears to be essential spawning and rearing habitat for summer-run steelhead in the Eel River. Along with the Upper Middle Fork of the Eel and Upper Mainstem Eel above the dam, the South Fork Van Duzen would be sensible areas to focus restoration efforts and increase cold water refugia with the future threat of warming from climate change (High et al., 2006), which could reduce the consistency of spring flows and abundance of cold water habitat. In addition to restoration work in or adjacent to the river channel, it would be advantageous to further encourage beneficial management of the South Fork Van Duzen watershed, both on the publicly and privately-owned lands, as well as the road and highway system, to improve holding, spawning and rearing habitat.

Dams and diversions are one of the primary causes of decline in salmonid populations (Katz et al., 2013). The phase in U.S. history of constructing new dams has generally ended (Graf, 1999) and some once dammed rivers are being returned to a free-flowing state through the FERC re-licensing process (Bednarek, 2001). The re-licensing of the project hydropower project that contains the two dams on the upper Eel and the potential for the return of anadromous fish presents questions regarding how best to restore salmonid runs to the upper basin. Our results suggest that after being isolated from anadromous runs of steelhead for nearly 100 years, the resident trout population above Scott Dam appears to have retained the potential to express both anadromy and premature migration, although we cannot estimate how long this diversity will continue to be maintained in the resident population. It has been shown that once premature migrating Chinook are extirpated from a system, premature migrating alleles are not maintained in the mature migrating population due to co-dominance at the *GREB1L* locus (Thompson et al., 2019). Heterozygotes have an intermediate phenotype (Prince et al., 2016, Thompson et al., 2019), therefore express an intermediate migration time, leading to decreased fitness (Thompson et al., 2019). However, it remains unclear what, if any, types of selective forces resident trout with the premature migration genotypes experience. It’s possible that the run-timing genetic variation may not lead to substantial phenotypic differences in resident trout populations, but further study into the degree of phenotypic differences experienced by resident trout carrying the premature or mature migration genotypes should be undertaken.

Dams have forced many populations of *O. mykiss*, which once were suffused with highly fecund individuals fattened on ocean nutrients, into being composed solely of smaller, resident individuals (Clemento et al., 2009). It has been shown that in some populations isolated above barriers the potential for anadromy is lost (Pearse et al., 2014), but in others it is retained (Holecek et al., 2012, Leitwein et al., 2016, Arostegui et al., 2019). Due to their propensity to conserve the ability to smolt, Phillis et al., 2016, concluded that an above barrier population of *O. mykiss*, isolated for around 100 years as well, could aid in the recovery of a related, proximally located anadromous population. Our results indicate that the resident trout population above Scott Dam could exhibit an anadromous life-history, given the opportunity. The genotype associated with anadromy has been mostly lost from the population isolated above Bloody Rock Roughs because trout that out migrate are likely unable to navigate back upstream through the roughs, an example of purifying selection observed elsewhere above barriers (Pearse et al., 2009). However, if anadromous fish were able to access that stretch, they could recolonize and re-introduce the anadromous genotype and, potentially, the already present premature migration genotype could eventually be utilized by these anadromous individuals. With no assistance from hatchery fish, summer-run steelhead quickly re-established in the Elwha River after dam removal (McMillan et al., 2019). It has been shown in other salmonid populations that retention of a diverse array of life-history strategies increases a population’s resiliency (Hilborn et al., 2003). Population structure analysis confirmed that *O. mykiss* above Scott Dam are of Eel River origin. Our results suggest that, considering their present state of run-timing genotypes, the potential to exhibit migratory behavior, and overall genetic diversity, the resident trout population above Scott Dam would be primed for re-establishment of steelhead post dam removal. Given the results of our study and the potential negative consequences and costs of hatchery fish, it seems prudent to give the native *O. mykiss* the opportunity to autonomously re-establish anadromy in the upper watershed upon dam removal.

Given the dramatic and continued decline of summer steelhead relative to their historical abundance, understanding if and to what extent the summer-run allele is maintained in the absence of the summer-run phenotype is a critical question for conservation and restoration efforts. We explored this issue in the South Fork Eel, where historical documentation describes summer-run steelhead, but the contemporary population appears to only contain winter-run individuals. We did not observe the summer-run allele in nearly 1600 samples, even though the samples were primarily collected from Elder and Fox Creeks, key spawning tributaries in the section of river where the historical documentation describes summer-run. The lack of evidence of summer-run alleles in this population is inconsistent with the idea that the summer-run variant is maintained as standing variation in populations that lack the summer-run phenotype (Waples & Lindley, 2018, Pearse et al., 2020). Thus, in contrast to what we observed above Scott Dam, the summer-run allele does not appear to persist in the absence of the summer-run phenotype in anadromous waters, even when a relatively healthy resident population is present as is the case in the upper South Fork Eel (Kelson et al., 2019, Kelson et al., 2020). Considering that the summer-run allele is necessary to express the summer-run phenotype (Prince et al., 2017), the absence of summer-run alleles in the South Fork Eel distinctly reinforces the need to protect the summer-run phenotype in the rest of the Eel River watershed and throughout Northern California (Waples & Lindley, 2018).

Currently, Eel River salmonids present one of the greatest opportunities for fisheries recovery on the West Coast of North America, with the lack of hatcheries, the hydropower project up for re-licensing, a broad-interest coalition working towards a compromise regarding the dam for out-of-basin water users and ESA-listed salmonids (Eel River Forum Members, 2016), no major urban populations in the watershed, reformation of historically damaging logging practices, and the persistence of four species of salmonids, with multiple independent populations (Yoshiyama and Moyle, 2010). The case for additional protections for summer-run steelhead in the Eel River has been made before, even with findings of less substantial genetic differentiation between the two populations (Nielsen and Fountain, 1999). Considering our results and the continued decline of summer-run steelhead in the Eel River and beyond, it is likely that current management approaches (e.g., NMFS, 2016, Pearse et al., 2020) are inadequate to protect these unique fish into perpetuity. However, the results from this and other recent work (e.g., Cooper, 2017, McMillan, 2019) present an opportunity to design conservation, management, and restoration strategies that would increase the potential for long-term persistence of summer-run steelhead in the Eel River and beyond.

## Materials and Methods

### Field Sampling

Sampling locations were selected to obtain a broad spatial distribution of the Van Duzen, Middle Fork and upper mainstem Eel Rivers with additional focus upstream and downstream of potential seasonal and anadromous barriers. Potential barriers were identified by a combination of historical documentation, communication with local biologists, and ten years of observations during annual dive count surveys by two of the authors. Spatial gaps remain in the sampling distribution in the lower Middle Fork Eel, Black Butte (tributary to Middle Fork Eel) and upper Van Duzen Rivers due to warm temperatures, lack of cover, and resultant low density of steelhead juveniles. We collected 834 upper caudal fin clip samples from young of the year (YOY) (age 0) (n=636), juvenile (age >1), (n=179), and adult anadromous steelhead (n=19) in the three basins from June 2016-November 2018. YOY were primarily sought due to their abundance, relative ease of collection, and being representative of adult spawning distributions (Hudy et al., 2010). The majority of sampling was conducted between July-September 2017 and July-September 2018 using hand-held dip nets. Hook and line sampling was used to capture larger residents and adults. Lastly, all carcasses encountered (n=13) were sampled opportunistically. Approximately ten to twenty samples per year were collected from each site and each fish’s fork length was recorded. Age class was determined based on length and date of capture, with a maximum size cutoff of 70mm in July and 110mm in September, to be called YOY. Within the YOY age class attempts were made to get individuals from a wide range of sizes and spread over longer spatial distances (hundreds of meters) to reduce sibling capture. Samples from the Upper South Fork Eel River in Angelo Reserve (mainstem n=6, Elder Creek n=1301, Fox Creek n=286) were collected following methods described in Kelson et al., (2020), using an electrofisher and hand-held dipnets from 2014-2017. Sampling was conducted under multiple years of NMFS 4D and CDFW Scientific Collection permits. Fin clips were placed on filter paper in small envelopes, dried, and stored at room temperature for later DNA extraction.

### Laboratory Processing

A 2-5mm^2^ piece of each sample was placed into individual wells with Lifton’s buffer in 96-well plates. Plates were stored at −20^°^C to await extraction. DNA was extracted using Ampure XP beads and protocol described by Ali et al. (2016). The extracted DNA was then used to prepare Rapture (RAD-Capture) libraries according to protocol in Ali et al. (2016), with 516 baits. 500 baits spread across the 29 chromosomes of the *O. mykiss* genome were designed by Ali et al. (2016), and the remaining 16 baits targeted RAD loci in the *GREB1L* region (Table S6). The baits were obtained from a MYcroarray MYbaits kit from Biodiscovery, LLC. The variable amounts of DNA in each well were standardized using an Eppendorf epMotion 5070. Each sample was given a unique combination of plate and well barcodes so that all samples could be combined for sequencing in a single reaction. We had two separate library preparation and sequencing runs, first in 2017 and then again in 2018 for the different years of sampling. The two sequencing runs were given 10% and 20% of an Illumina HiSeq 4000 lane, respectively. Samples from 2017 that had low read counts (n=152) were re-sequenced with the 2018 samples and reads were merged using SAMtools merge.

### Population Structure and F_ST_

After demultiplexing (Ali et al., 2016), samples were aligned to a recently assembled rainbow trout reference genome (Pearse et al., 2019) using BWA-MEM (Li and Durbin, 2009), then filtered to remove non-properly paired reads and PCR duplicates using SAMtools view and rmdup. Only samples with greater than 5000 filtered alignments (n=761 combined for the Middle Fork, Van Duzen and upper main Eel) were retained for population structure analyses. In addition, only YOY samples were used for analyses discussed in this paper. We removed putative large family groups identified through Principle Component Analysis, which dominated PC1, from two sites, SEE and URS (Table S1). Beyond that we did not remove any individuals based on family structure due to the numerous potential consequences and lack of explicit benefit to our analyses in doing so (Waples and Anderson, 2017). Larger juveniles and adult samples confirmed previous association with G1L genotype and phenotype (e.g., the adult samples we collected in July and August in the Van Duzen (n=9) and Middle Fork (n=5) were found to be homozygous for the summer-run genotype) (Prince et al., 2017) (Table S1). There are two notable exceptions to the practice of only including YOY in our analyses, in the case of locations in the upper mainstem Eel, above Scott Dam, where there are no anadromous fish and densities of fry were low. Resident samples were included from this basin as they are presumed to not migrate between the two sampling regions (Rice Fork and upper mainstem) due to the Bloody Rock barrier. The other exception was in the upper Middle Fork, also above anadromous barriers, where YOY densities were too low to conduct analyses without using juvenile samples. The barriers in this region also preclude juveniles spawned elsewhere from immigrating.

Analysis of Next Generation Sequencing Data (ANGSD) software (Korneliussen et al., 2014) was used to generate covariance matrices for principal component analysis (PCA). The PCA was run, following methods from Prince et al., 2017, with slight variations, by identifying polymorphic sites with a SNP_pval of 1e-12, determining major and minor alleles, doMajorMinor 1, estimating allele frequencies, doMaf 2, and retaining SNPs with a minor allele frequency of at least 0.05. The PCA was run on all sequenced loci, excluding OMY5 which contains a large number of SNPs in strong linkage disequilibrium and can skew PCA results. R software (ggplot2, Wickham, 2016) was used for visualization of PCA results. Pairwise F_ST_ between sites were calculated by using ANGSD to create a site frequency spectrum with the command doSaf. RealSFS (ANGSD) was then used to calculate a two-dimensional site frequency spectrum and global F_ST_ for each pairwise combination. Sites with less than five samples were combined with neighboring locations for a total of 42 sites and 861 pairwise site combinations. Distances between sites, in river kilometers, were calculated using the R package riverdist (Tyers, 2017). Pairwise combinations were then grouped by presence of a barrier between locations and F_ST_ values were scaled (F_ST_ /1-F_ST_). Plots were created using the R package ggplot2 (Wickham, 2016). Only the pairwise site combinations in the Van Duzen River are referred to in this paper (sites=27, pairwise=276). Significance of difference in pairwise F_ST_ values around barriers was tested by randomly assigning sites to be above or below a barrier and calculating the difference in the y-intercepts of site combinations that do not cross a barrier to those that cross the Eaton Falls and South Yager Barriers, separately. The y-intercept represents the expected Fst between two sampling locations directly adjacent to each other (i.e., zero river kilometers). This random assignment was replicated 1000 times in R to see how frequently the randomized difference in y-intercepts was greater than or equal to the observed difference in y-intercepts. If a potential flow dependent barrier inhibits migration, the y-intercept from the site pairs that cross this barrier should be significantly greater than the y-intercept from site pairs that do not cross a barrier, and the extent of this difference should relate to the extent to which the barrier inhibits migration.

### Genotype Calling and Genetic Diversity

Genotype calls at the *GREB1L* region were made using two SNPs on OMY28, at positions 11667773 and 11667954, that are on different RAD loci from the same SbfI site. We used ANGSD (Korneliussen et al., 2014) -doGenos to determine genotype likelihoods at each of two SNPs. We then calculated the combined likelihood of the three possible genotypes for each individual by taking the product of the likelihoods from each SNP. We then calculated genotype posteriors using the combined likelihoods assuming a uniform prior and used a posterior cutoff of 0.8 (n=661 combined for the Middle Fork, Van Duzen and upper main Eel and n=1593 for the South Fork Eel) to call the run-timing genotypes, winter-run, summer-run, and heterozygotes.

Genotype calls at the OMY5 region were made by creating a covariance matrix using ANGSD of the 19 RAD tags sequenced on the OMY5 chromosome (Kelson et al., 2019). This generated three distinct groupings of PC1 scores, −0.045- −0.03, −0.006-0.015, and 0.045-0.06, that correspond to the “resident”, “heterozygote” and “anadromous” genotypes (Figure S1; Kelson et al, 2019), respectively. Outliers were not included in genotype frequency calculations (n=48) (Supplemental Figure 1).

Maps of genotype frequencies were made using R packages ggplot2 (Wickham, 2016), scatterpie and ggmap (Kahle and Wickham, 2013). Per base average pairwise nucleotide differences (π) values were calculated with ANGSD by using realSFS to create site frequency spectrums for each location. These values were grouped by basin and the mean values were calculated for each basin. Significance between means was tested using the multcomp package in R (Hothorn et al., 2008).

## Acknowledgements

We thank J. Abrams, E. Bertz, G. Bramble and P. Bramble, M. Clapp, J. Crawford, R. Dana, S. Harris, D. Heaton, M. Heaton, W. Heaton, L. Holm, C. Holmgren, K. Lackey, D. and M. Moore, N. Okun, B. Pagliuco, S. Ricker, S. Rizza, A. Saley, E. Stockwell, R. Thompson, S. Thompson, T. Thompson and L. Woodcock for assistance with access and sample acquisition. We also thank L. Holm, R. Peek and S. Rizza for assistance with map and data visualization. And we thank E. Habibi, A. Rypel, A. Schreier and T. Thompson for invaluable insight and feedback on earlier versions of this manuscript. We would like to recognize E. Bertz and C. Holmgren for their exploratory efforts which led to the re-discovery of the Van Duzen River summer-run steelhead population and their contribution to the nomenclature of features in the Lost Duzen region. And finally, we commend S. Harris and S. Thompson for their unwavering commitment to the study and preservation of summer-run steelhead in the Eel River.

---

**Supplemental Figure 1.**
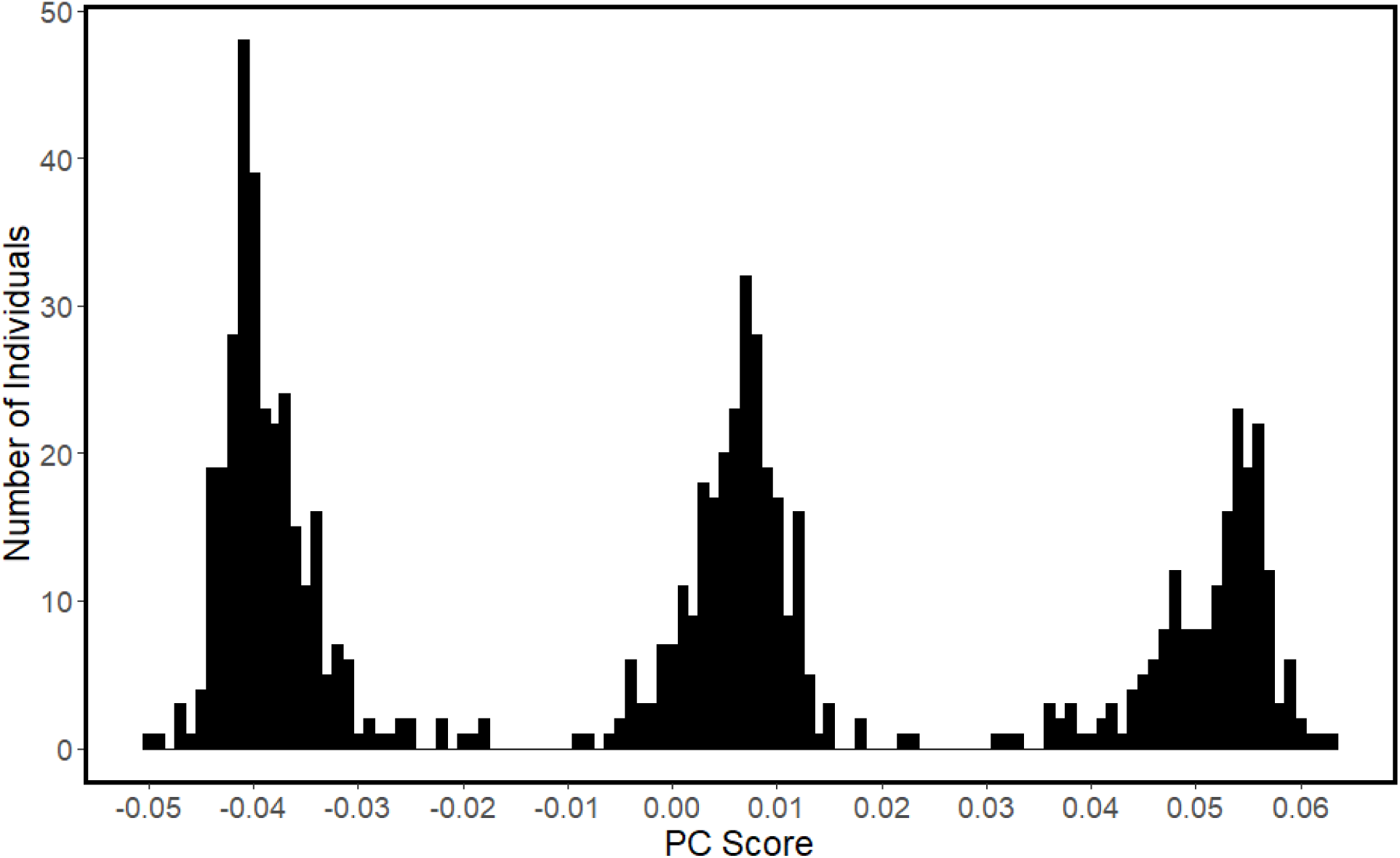
Bar plot showing distribution of Principal Component 1 values on chromosome 5 or OMY5. Three clear peaks indicate resident, heterozygous and anadromous genotype, from left to right.

